# TimeTraits: extracting functional traits from biological time-series in R

**DOI:** 10.64898/2026.02.20.706990

**Authors:** Sarah C. L. Lock, Marina I. Knight, Seth J. Davis, Daphne Ezer

## Abstract

**Summary:** TimeTraits provides a series of functions to enable the extraction of parameters from time series data using Functional Data Analysis methods. We demonstrate its utility in dissecting the changes in curve shape of a circadian clock bioluminescence marker, following a photoperiod shift in wildtype and *phyB* Arabidopsis.

**Availability and Implementation:** TimeTraits is available in CRAN: https://cran.r-project.org/web/packages/TimeTraits/readme/README.html

## 1. Introduction

Circadian rhythms govern essential processes in animals and plants, including sleep-wake cycles, photosynthesis and growth. For animals, understanding circadian rhythms is crucial for timing medical interventions (Dibner and Schibler 2015; Ayyar and Sukumaran 2021). In plants, they offer insights into optimising agricultural strategies and enhancing resilience to environmental stressors (Ullah et al. 2024; Schmidt et al. 2024; Ficht et al. 2023; Minoli et al. 2022). Therefore, how these rhythms are analysed is very important.

Traditionally, the Fast Fourier Transform and Non-linear Least Squares (FFT-NLLS) approach has been widely employed for analysing bioluminescence circadian data (Brancaccio et al. 2019; Gould et al. 2017; Moore, Zielinski, and Millar 2014). However, this method assumes that the underlying rhythms are stationary (i.e. that the rhythms are consistent over the time series) potentially leading to oversimplified interpretations and limiting the ability of scientists to investigate rhythms under environmental conditions that change over time. It has been shown that expression of core *Arabidopsis thaliana* clock genes over time do not look like typical sine-waves and are in fact non-stationary (Hargreaves et al. 2016, 2019). Alongside this, often biologists summarise properties of the circadian clock in terms of three key features: amplitude, phase, period. By exclusively investigating these features, information about how the gene expression curves deviate from sine-waves are overlooked.

Functional Data Analysis (FDA) statistical methods consider the entire profile of gene expression over time, without many assumptions about which features are important (Levitin et al. 2007; Ramsay, Hooker, and Graves 2009). This approach offers a more nuanced and flexible framework for analysing time-dependent data, particularly suitable for circadian data analysis. Viewing data as functions rather than as discrete points, enables the exploration of dynamic patterns including subtle changes in waveform and any irregularities happening over time. This approach gives the ability to uncover hidden patterns and complexities that might be overlooked by conventional methods. Furthermore, FDA techniques offer powerful tools for functional regression, clustering, and classification, allowing for the integration of covariates and the identification of distinct temporal patterns within datasets (Ramsay et al. 2024; Ramsay, Hooker, and Graves 2009). This capability is particularly valuable in circadian research, where external factors such as light exposure, genetic variations, or environmental conditions can influence rhythmic patterns. By incorporating these additional features into the analysis, FDA facilitates a more thorough investigation of the underlying mechanisms governing circadian rhythms and their interactions with external stimuli.

The adoption of FDA approaches for circadian data analysis has yet to be extensively explored due to the lack of accessible implementation. While R packages exist that implement FDA strategies, data wrangling is required to appropriately format data from lab hardware, and the analysis workflow involves multiple steps with many decision-points that a biologist may not be well suited to make. To address this gap and promote the utilisation of FDA techniques within the circadian community, our goal is to develop an R package specifically tailored for this purpose. By providing user-friendly functionalities for data preprocessing, curve estimation, functional principal component analysis and visualisation, our package aims to lower the barriers to facilitate the exploration of the dynamic nature of circadian rhythms.

### 2. Methods

The ‘package’ has been published at the Comprehensive R Archive Network (CRAN). It is designed to aid the exploration of circadian luminescence data, by providing the user with a collection of prespecified analysis steps thus reducing the complexity of applying FDA methods. The package offers three core functions—(i) filter_curves: for pre-processing data directly from bioluminescence lab hardware and performing data quality controls and (ii) smooth_fun: for turning the rhythmic data into normalised functions and (optionally) filtering potential outlier curves. (iii) functional_traits: for extracting a set of non-stationary properties of the rhythms using FDA approaches.

The package conducts three main functions for analysis, starting with preprocessing of the data. The provided R function filter_curves aims to filter circadian luminescent data based on specified criteria, preparing it for further analysis. Initially, the function validates the input data is in the correct format, next a data frame containing curves within the specified time range (`from` to `to`) is created along with the corresponding time vector. Only curves exceeding the minimum luminescence threshold (`min_lum`) are identified and retained (SupFig 1).

Many researchers use luciferase assays to produce rhythmic bioluminescent readings, which have signals that decay over time. Moreover, some organisms may die over the course of the experiment and therefore lose their rhythmicity (Paajanen, Lane de Barros Dantas, and Dodd 2021; Creux and Harmer 2019). Filter_curves offers optional methods for de-trending the curves and filtering out samples that lose their rhythmicity in the last 48 hours (or an alternative value set by the user). Detrending is achieved by fitting a linear regression to each circadian rhythm over time and using the residuals to remove any trend (SupFig 2). A Lomb-Scargle periodogram analysis is performed for the last 48 hours of the analysis to identify weakly rhythmic curves (SupFig 3) (Ruf 1999). Curves failing to meet the significance threshold (0.001) in the periodogram analysis are removed from the dataset. Finally, the filtered curves are combined back with the corresponding time vector into a final data frame, which is returned as the output of the function, ready for further analysis.

The next function the package provides to the user is aimed to estimate the discrete curves into smooth continuous curves using FDA methods (`smooth_fun`). Initially, the function checks if the input data is correctly formatted, typically this input will be the preprocessing output. Subsequently, each luminescent curve is normalized to the range [0, 1]. The function then proceeds to create a list containing each curve along with its corresponding time values, removing any missing data points. For each curve, basis functions and functional parameters are generated using FDA methods (Ramsay et al. 2024). This involves defining a basis function via B-splines and setting a penalty for roughness that controls the smoothness of the fitted function. The user may define these parameters manually or they can be optimally selected using the helper function `select_basis_lambda_LOOCV.` This function performs leave-one-out cross-validation across a grid of basis numbers and smoothing parameters (λ), identifying the combination that minimises error (SupFig 4). `smooth_function` then applies smoothing to each curve individually (using the chosen basis number and λ), producing smooth basis functions. Next, a new time vector is created to cover the range of all original time values. The curves are evaluated under this new time vector, resulting in a matrix of all curves under the same time points. Outliers in this dataframe are detected using FDA outlier detection methods (‘fdaoutlier’), specifically the Total Variation (TV) Smoothing and Derivative-based Measures of Scale (MSS) (Febrero–Bande and Fuente 2012). Detected outliers are then optionally removed from the dataframe. Finally, the function returns the smoothed curves across the desired time frame providing users with functional data objects ready for further analysis.

The final function in this package (`functional_traits)`, performs functional principal component analysis (FPCA) on smoothed circadian time-series data and extracts group-level traits from the resulting FPCA score space. The analysis can optionally be split by time segments (e.g. pre/post environmental shift) and by derivatives of the curves. When segments are supplied, functional observations from all segments are first re-evaluated onto a shared time grid and analysed jointly, ensuring that FPCA scores are directly comparable across segments. For each derivative, FPCA is performed once across all samples and segments simultaneously. Segment- and group-specific traits are then computed post hoc from the shared FPCA score space. The traits are outputted in a table compatible with further R/qtl analysis. Traits calculated include mean and standard deviation of FPCA scores (per principal component), mean and standard deviation of distance to the FPCA-space centroid and convex hull area in PC1–PC2 space (measure of within-group diversity).

## 3. Example

Most circadian research in plants has focused on constant conditions over time, because of the limitations of the analysis approaches. Here we show that our approach enables us to investigate rhythms under environmental conditions that change over time, specifically having shifts in day lengths in the middle of the experiment, which could be applied to investigate how plants respond to the change in seasons (Ronald et al. 2024; Lock et al. 2025; Mehta et al. 2025). The package comes with this data set manually compiled (phyb). It contains samples of luminescence data from *Arabadopsis thaliana* seedlings from Ws-2 (harbouring CCR2:LUC) and then Ws-2 plants containing a mutation in phytochromeB gene (*phyB-10, also* harbouring CCR2:LUC).

Seeds were surface-sterilized and plated onto MS medium with 3% sucrose and then stratified for 4 days. After stratification, seedlings were entrained under a short day (SD, 8 hours light: 16 hours dark) photoperiod of cycles with a constant temperature of 22° for 7 days. On day 6, seedlings were transferred to black 96-well Microplates with MS medium containing 3% sucrose. Plants were superficially treated with 15 μl 5 mM D-Luciferin. Seedlings were then re-entrained for 1 day under the respective entrainment conditions before being transferred to the TOPCOUNT (Perkin-Elmer [Perkin-Elmer-Cetus], Norwalk, CT) where luminescence measurements began. On day 11 seedlings were then shifted into a long day (LD, 16 hours light: 8 hours dark) photoperiod (see Figure 1A). TOPCOUNT experiments were carried out under blue–red light and a constant temperature of 21°.

**Figure 1:**
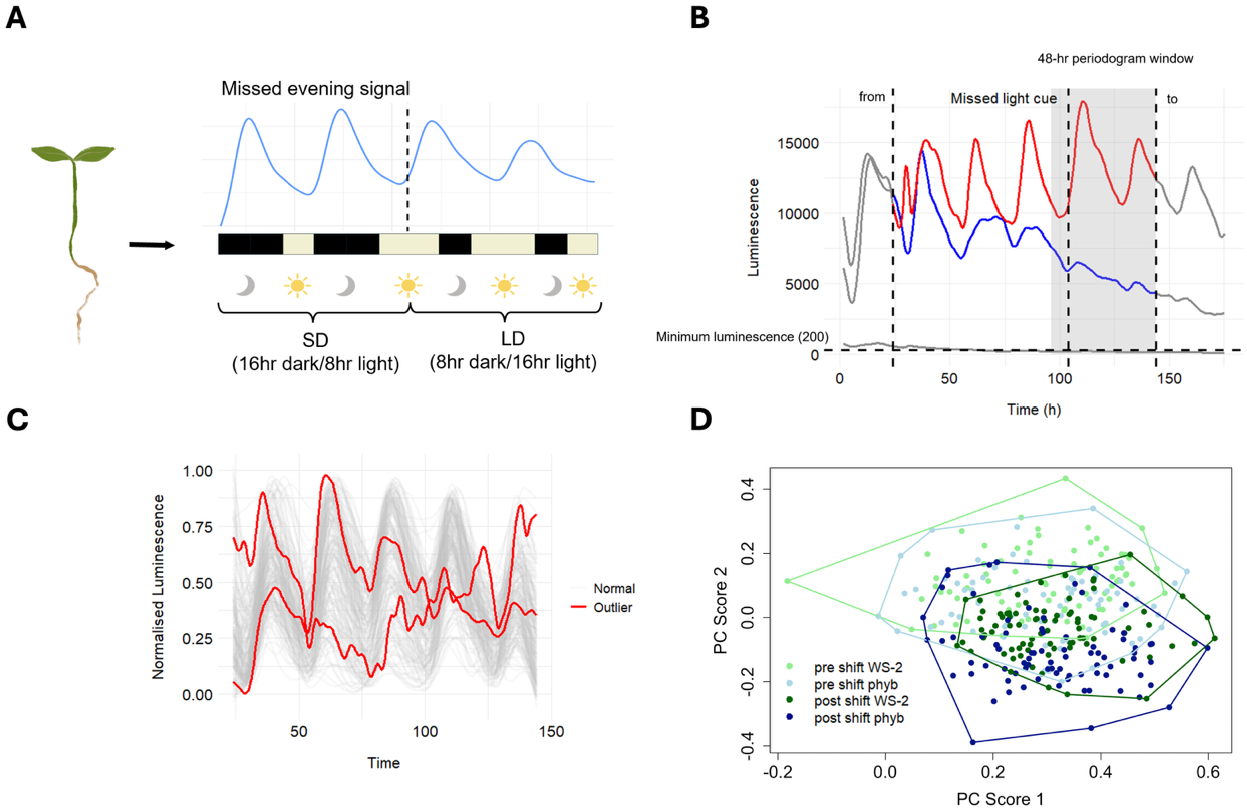
Overview of the TimeTraits analysis pipeline using circadian luciferase data. **(A)**. Experimental design. *Arabidopsis thaliana* seedlings (Ws-2 wild type and phyB mutants) expressing a luciferase reporter were entrained under a short day (SD, 8 hours light: 16 hours dark) and subsequently shifted to a long day (LD, 16 hours light: 8 hours dark) while luminescence of CCR2:LUC reporter gene was continuously recorded. **(B)** Illustration of curve filtering using the filter_curves function. Shown are the user-defined time window (“from”, “to”), minimum luminescence threshold, and the 48-hour periodogram window used to assess rhythmicity. Grey curves indicate traces removed during filtering. The blue curve highlights an example rejected due to lack of rhythmicity in the selected periodogram window, while the red curve represents a trace that passed both the minimum luminescence and rhythmicity criteria. **(C)** Normalised and smoothed luminescence profiles produced by the smooth_fun function. Grey curves represent retained samples, while red curves indicate outliers removed based on overall shape characteristics. **(D)** Functional principal component analysis (FPCA) of smoothed curves, showing PC1 versus PC2 scores for pre and post photoperiod shift segments across genotypes. Convex hulls summarise group-level structure in FPCA space.

The `filter_curves` and the `smooth_fun` functions were used to perform filtering for unsuitable rhythms and further analysis. The minimum threshold was set at 200 and looking at a window of 44 -144 hours and a 48-hour periodogram window, Figure 1B shows examples of curves that would be included and excluded under these selection parameters. Clear smooth functions were outputted after the removal of outlier curves based on shape (Figure 1C). After acquiring smooth curves these plants were then evaluated as pre and post photoperiod shift and a FPCA applied using `functional_traits` (Figure 1D). Our analysis shows that our methods effectively filter arhythmic genes, remove outlier curves, and produce smooth normalised curves. These were then used to perform downstream analysis that could not be performed under FFT-NLL, such as producing an FPCA plot that shows how the curves change after a photoperiod shift denoted by a clear separation in pre and post shift plants (Figure 1D).

## 4. Conclusion

In conclusion this package successfully achieves its aim of providing an accessible and automated workflow for analysing circadian time series data using FDA approaches. By integrating the processing, filtering, detrending, outlier removal and functional smoothing into a single framework it reduces the complexity of applying FDA methods. Compared with traditional approaches to analyse circadian data which assume stationary and focus on a small number of rhythmic parameters this package captures the full functional profile of circadian curves enabling the exploration of non-stationarity and response to external cues over time. This advancement broadens the analytical possibilities for circadian researchers, supporting biologically meaningful exploration and interpretations of rhythmic data.

## Acknowledgements

We thank Jason Daff, Paul Scott, Harry Stevens and the rest of the University of York Horticulture team. The Viking Cluster was used in this project, which is a high-performance compute facility provided by the University of York. The authors are grateful for computational support from the University of York High Performance Computing service, Viking and the Research Computing team. We also thank Katie Kindleysides for her graphic design of Arabidopsis images.

## Author contributions

S.C.L.L.: conceptualisation, methodology, data collection, software, analysis, investigation, writing—original draft, and visualization. M.I.K: conceptualization, methodology, writing—review and editing, supervision, and funding acquisition. S.J.D.: conceptualization, writing—review and editing, supervision, and funding acquisition. D.E.: conceptualization, methodology, analysis, writing—review and editing, supervision, and funding acquisition.

## Supplementary Information

Supplementary Information: See Supplementary document

## Data availability

TimeTraits is an open-source CRAN R package. Source code and example datasets used in this study are available at https://github.com/scllock/TimeTraits.

