## Supplemental Figures for "TimeTraits: extracting functional traits from biological time-series in R"

**Supplementary Information**

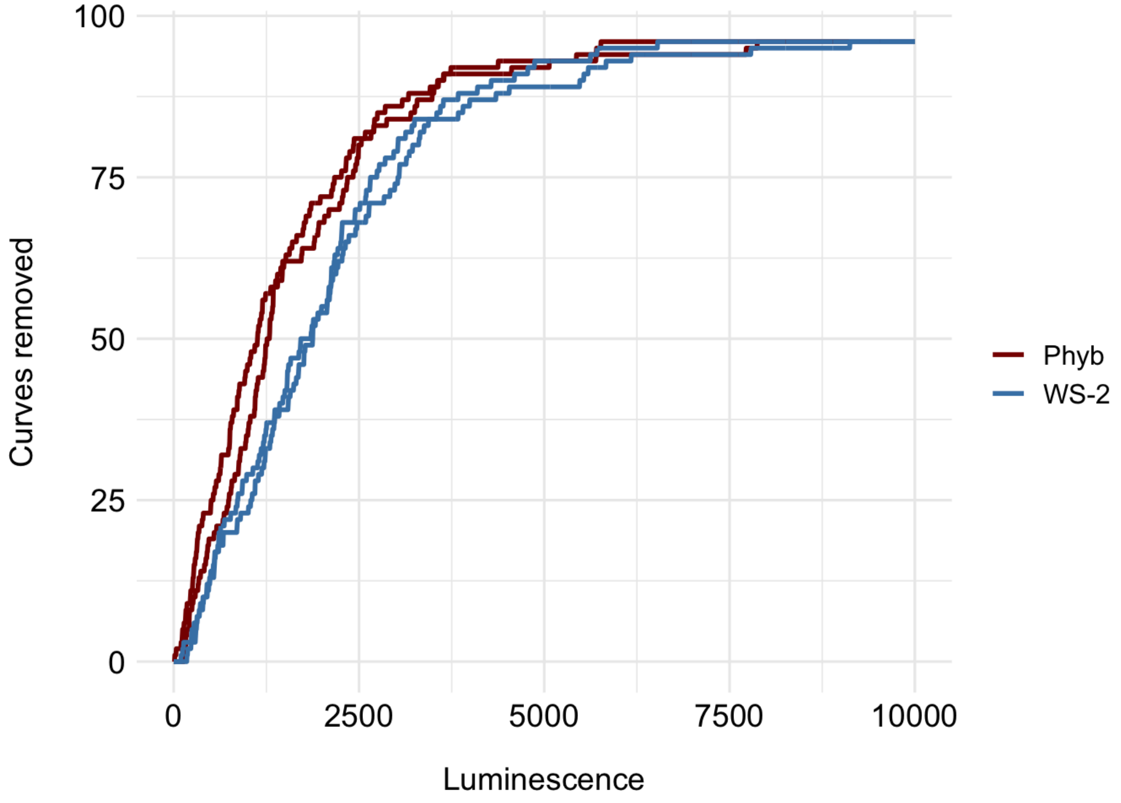


**Supplementary Figure 1:** Shows the number of curves removed within each genotype when increasing minimum luminescence threshold (`min_lum`) when using `filter_curves` function.

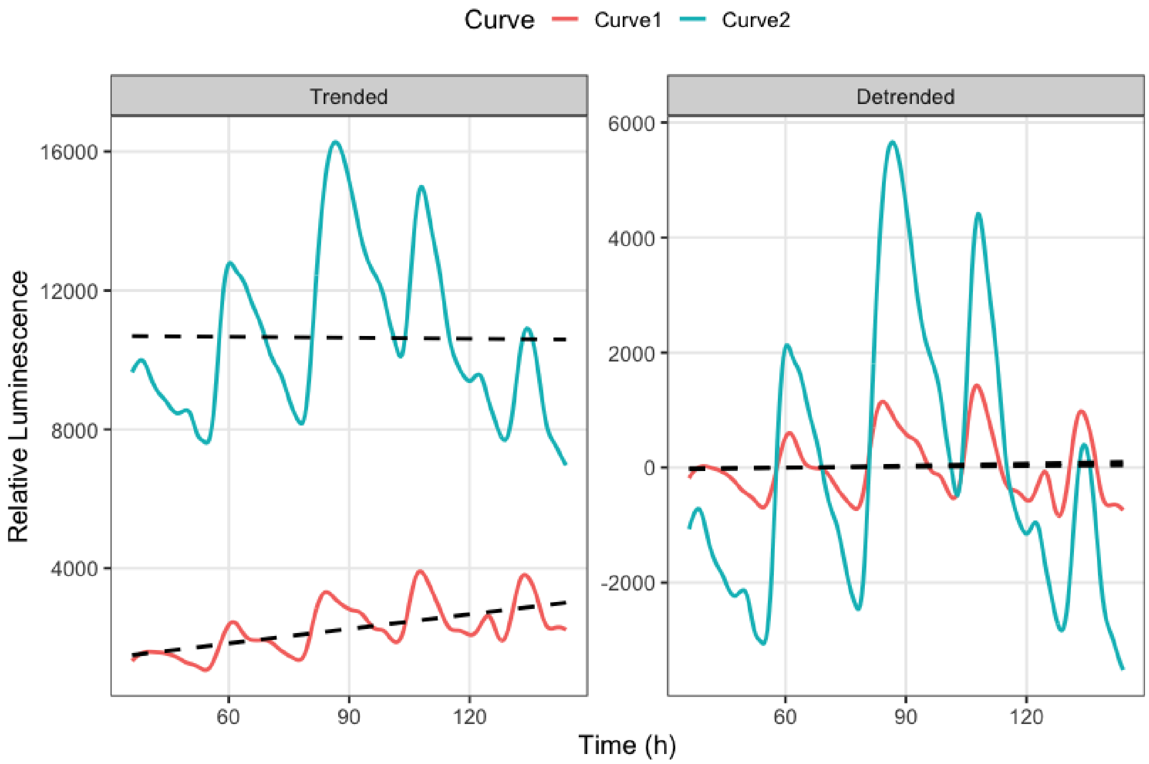


**Supplementary Figure 2:** Shows the effect of detrending the data when using `filter_curves` in two different example curves.

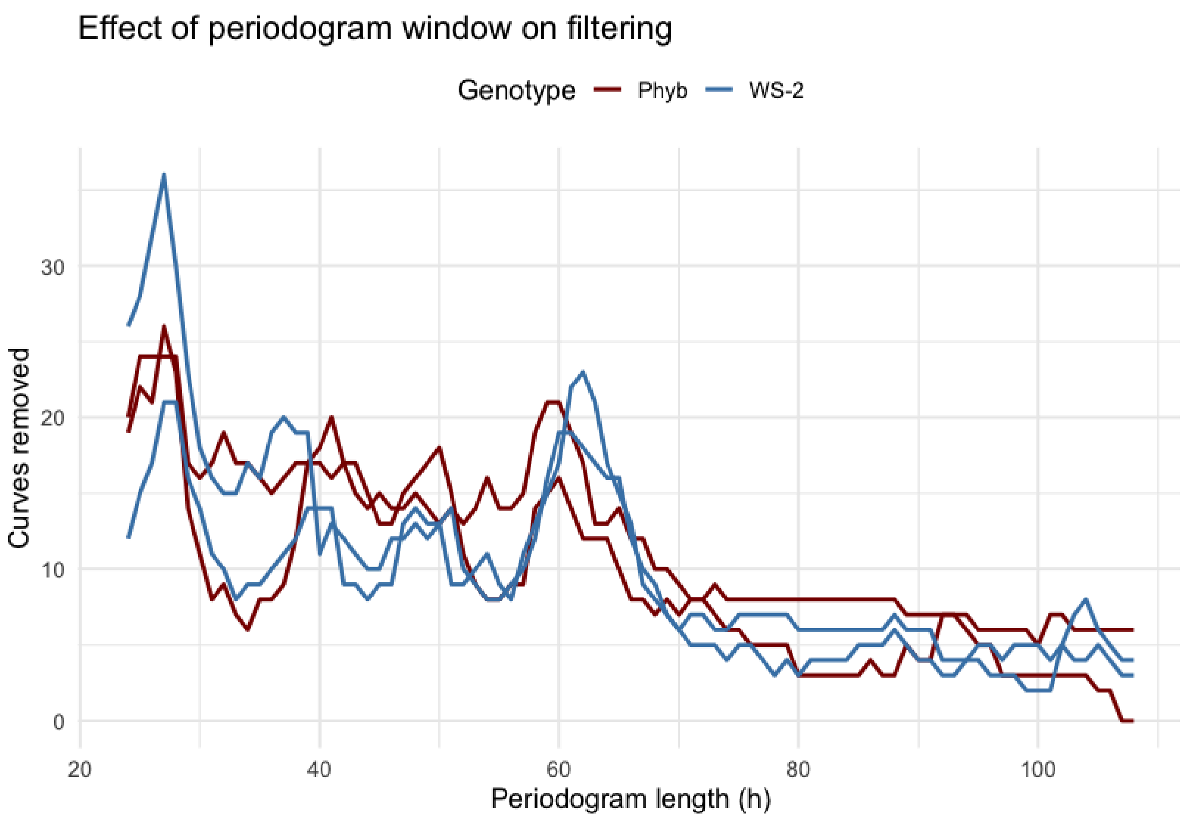


**Supplementary Figure 3:** Shows the effect of changing the length of the periodogram during filtering using `filter_curves`.

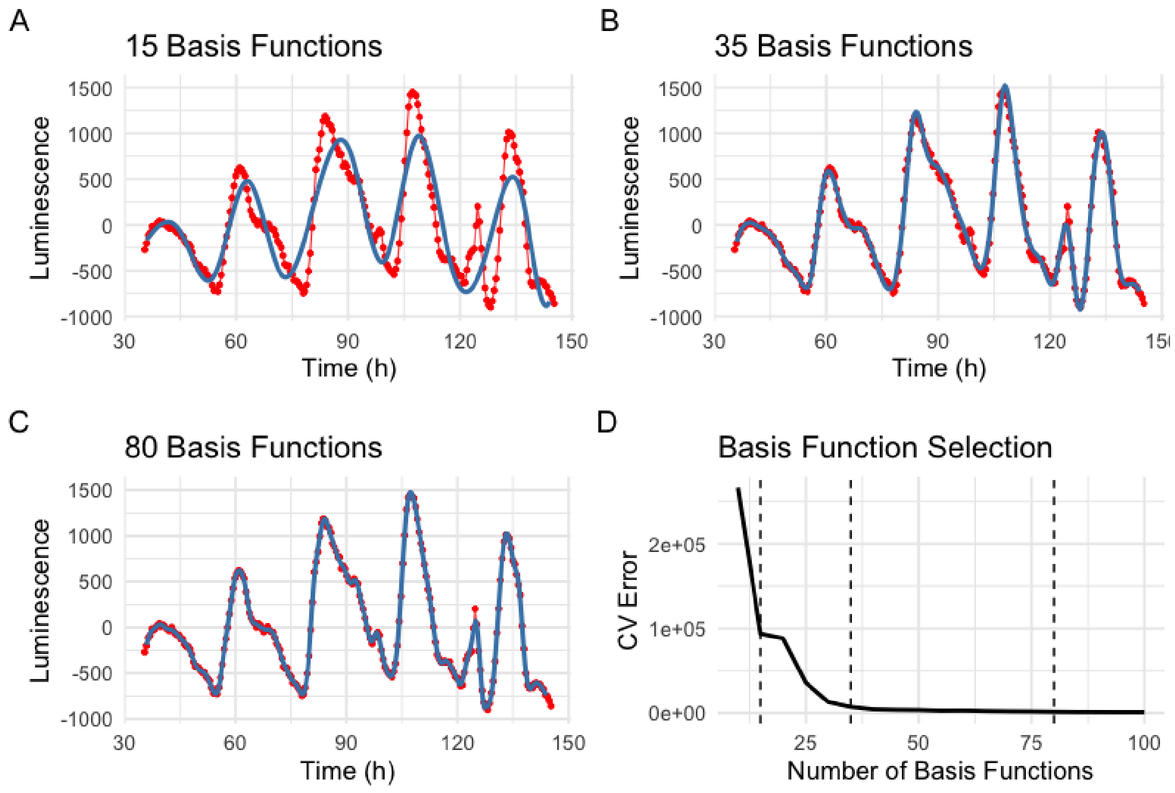


**Supplementary Figure 4:** Algorithm for picking number of basis functions and smoothing penalty using `select_basis_lambda_LOOCV`. (A – D) Shows an increase in basis functions decrease cross validation (CV) error. This reaches a plateau where more added basis functions does not improve fit and can lead to overfitting problems.
